# scPathoQuant: A tool for efficient alignment and quantification of pathogen sequence reads from 10x single cell sequencing data sets

**DOI:** 10.1101/2023.07.21.549987

**Authors:** Leanne S. Whitmore, Jennifer Tisoncik-Go, Michael Gale

## Abstract

Currently there is a lack of efficient computational pipelines/tools for conducting simultaneous genome mapping of pathogen-derived and host reads from single cell RNA sequencing (scRNAseq) output from pathogen-infected cells. Contemporary options include processes involving multiple steps and/or running multiple computational tools, increasing user operations time. To address the need for new tools to directly map and quantify pathogen and host sequence reads from within an infected cell from scRNAseq data sets in a single operation, we have built a python package, called scPathoQuant. scPathoQuant extracts sequences that were not aligned to the primary host genome, maps them to a pathogen genome of interest, here as demonstrated for viral pathogens, quantifies total reads mapping to the entire pathogen, quantifies reads mapping to individual pathogen genes, and finally reintegrates pathogen sequence counts into matrix files that are used by standard single cell pipelines for downstream analyses with only one command. We demonstrate that scPathoQuant provides a scRNAseq viral and host genome-wide sequence read abundance analysis that can differentiate and define multiple viruses in a single sample scRNAseq output.

## Introduction

Defining microbe/virus-host interactions of pathogen infections is reliant on understanding the effect of the agent on host cell gene expression concomitant with evaluating the dynamics of microbe/virus gene expression. For example, the emergence of SARS-CoV-2 and the COVID-19 pandemic underscore this need for integrated analysis of host and viral gene expression in single cells during acute virus infection. To facilitate understanding of host-pathogen interactions, experimental cell-lines and/or animals are typically experimentally infected with a known pathogen allowing this relationship to be interrogated. Integrated analysis of host and microbe/viral (referred herein as “pathogen”) gene expression allows for the discovery of molecular mechanisms of pathogen and/or host-mediated disease, and can reveal possible genetic targets for therapeutic intervention of infection. Similarly, such analyses are paramount for understanding the outcome of chronic virus infection such as hepatitis B virus or human immunodeficiency virus (HIV), and further demonstrate the need for a rapid and efficient analysis pipeline for quantifying genome-wide pathogen and host gene expression in single cell RNA sequencing (scRNAseq) experiments. The current process for quantification of pathogen RNA sequence reads with 10x single cell technology involves the following steps using CellRanger (Zheng et al. 2017): 1) integrating the pathogen genome into the host genome, regenerating the CellRanger genome index files, and then running the read alignment process or 2) performing a series of post-processing steps to extract unmapped reads from CellRanger output and aligning the extracted reads to the pathogen genome of interest. Both options have shortcomings. Option 1 requires more user steps as the target genome must integrated into the host genome and regenerated followed by read mapping, whereas Option 2 is highly cumbersome to require the implementation of several additional bioinformatics tools while additionally increasing the analysis time line. Additionally, 3 pathogen detection tools have also recently been developed including ViralTrack (Bost et al. 2020), PathogenTrack (Zhang et al. 2022) and Venus (Lee et al. 2022) for detection of viral reads in single cell data. These tools, while effective, require a number of preprocessing steps prior to pathogen alignment and read quantification. Moreover, these steps require a remapping of all reads to a host genome through STAR solo(Dobin et al. 2013) instead of pulling from already processed data for which CellRanger is often utilized.

Here, we report a new tool for alignment and quantification of pathogen sequence reads from 10x single cell sequencing data of known virus/microbe-infected cells. Our new tool, scPathoQuant (SPQ), minimizes the workload on users by packaging the necessary bioinformatics tools into an easy to install and use python library that can be executed with a single command. SPQ executes several analysis steps, including: 1) isolation of unaligned reads, 2) mapping of the isolated reads to the pathogen genome of interest with either bbmap (Bushnell 17 March 2014) or bowtie2 (Langmead and Salzberg 2012), 3) quantification of the resulting unique molecular identifier (UMI) read counts with HTseq (Anders, Pyl and Huber 2015) and 4) integration of the data set output back into the matrix files from the 10x scRNAseq data set. This integration step then allows the user to proceed with the analysis using common tools, such as 10x Loupe Browser (https://www.10xgenomics.com/products/loupe-browser), Seurat(Hao et al. 2021) or Monocle (Qiu et al. 2017), or any other standard scRNAseq analysis pipeline. Here we demonstrate the application of scPathoQuant for defining viral reads from single infection and multiple virus infection sample output from scRNAseq analyses.

### Design and Implementation

SPQ, initiates by extracting unaligned sequence reads from the host genome using SAMtools (Li et al. 2009), followed by alignment of read to the pathogen genome of interest using either bbmap (default) (Bushnell 17 March 2014) or bowtie2 (Langmead and Salzberg 2012) aligners (Fig 1). These two aligners were chosen for their speed and accuracy. BBmap and bowtie2 use different alignment algorithms, thus enabling the user to choose the aligner appropriate for their analysis (Table 1). Default parameters, such as rate of mismatches, insertions, and deletions (INDELs) vary across both alignment tools, resulting in slightly different outputs. SPQ uses bbmap and bowtie2 default settings, but the user may change any parameter of the aligners by adjusting the --bbmap_params or --bowtie2_params options, giving the user more control over the viral alignment process compared to CellRanger where individual STAR alignment parameters cannot be altered. UMI counts are quantified using HTSeq (Anders, Pyl and Huber 2015) first, counting any reads mapping to the pathogen genome (viral genome in our examples shown) and then quantifying individual gene counts if a gene transfer format (gtf) file for the specific pathogen genome is provided by the user. Quantifying UMI counts both for the whole pathogen genome and individual genes provides output to determine if pathogen transcripts exist in the cell while providing specific information on pathogen gene expression levels in each cell. UMI counts per cell are then integrated into the CellRanger raw and filtered feature matrix files (features.tsv.gz and matrix.mtx.gz) to allow for easy downstream analysis of pathogen abundance and specific gene expression levels concomitant with host gene expression levels. In addition, SPQ provides other output useful to the user including viral UMI count results for each cell in a csv file format, violin plots showing the distribution of pathogen UMI counts and coverage maps of the pathogen-mapped reads (Fig 1). Furthermore, SPQ generates a number of files that allow further in-depth analysis including bam files of reads mapping to pathogens of interest that can be easily uploaded into genome viewers for splice junction and SNP analysis in the sample, fastq files of all unmapped reads to the host, and fastq files of mapped reads to the pathogens of interest which can be utilized for phylogenetic analyses via blast or genbank.

**Table 1:** Alignment tools used in the analyses.

| Aligners | Alignment Version | Sequencing Data Type | Alignment Algorithm | Splice Aware |
| --- | --- | --- | --- | --- |
| *SPQ/BBMAP* | 38.9 | RNA/DNA | Multi kmer seed and extend<br>(http://bib.irb.hr/datoteka/773708.Josip_Maric_diplomski.pdf) | Y |
| *SPQ/Bowtie2* | 2.4.2 | RNA/DNA | Burrows-wheeler transform<br>https://genomebiology.biomedcentral.com/articles/10.1186/gb-2009-10-3-r25 | N |
| *CellRanger/STAR* | 2.7.2a | RNA | Maximal Mappable Prefix (*MMP*)<br>https://www.ncbi.nlm.nih.gov/pmc/articles/PMC3530905/ | Y |

**Figure 1.**
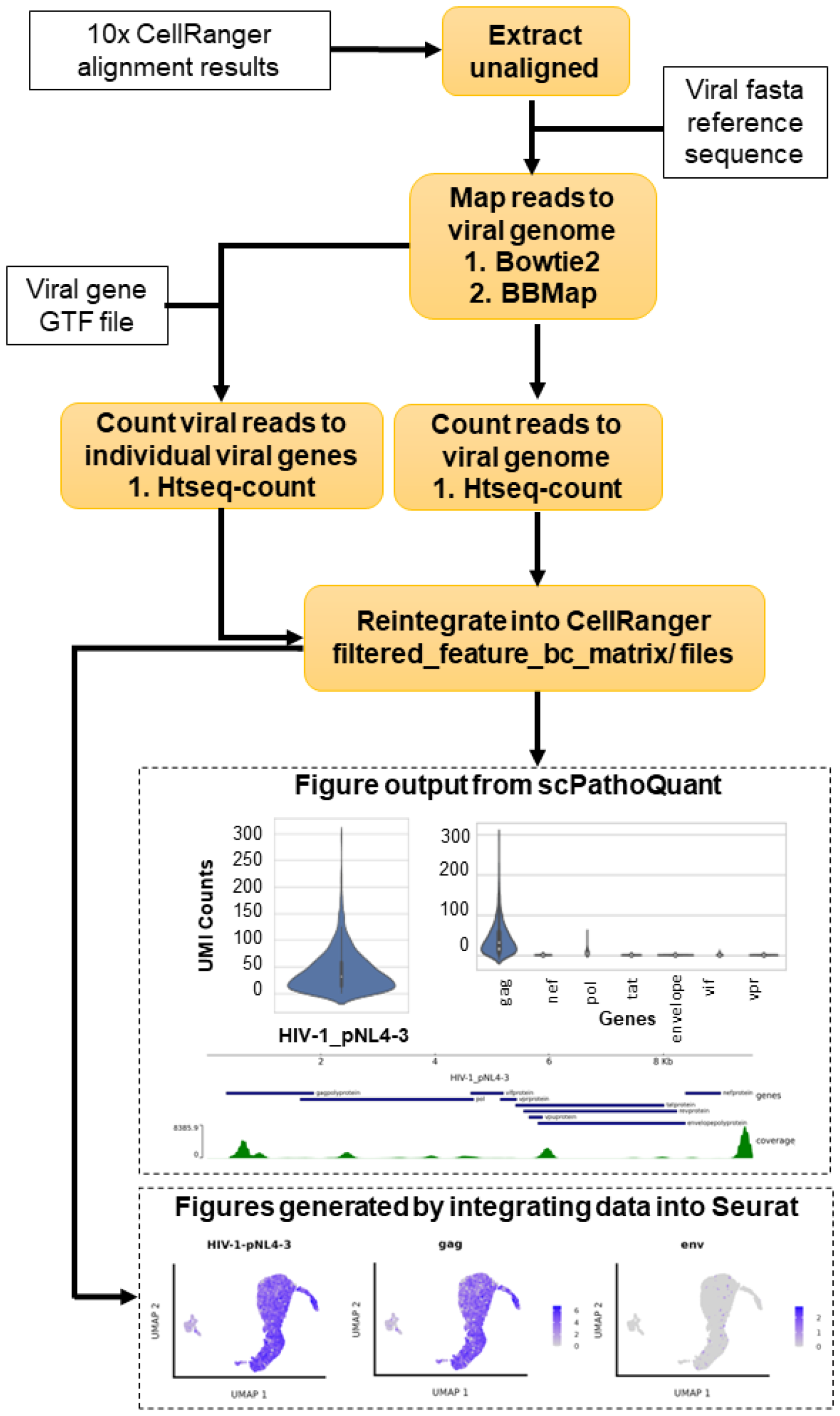
Software layout. Yellow boxes show the progressive steps implemented by SPQ analysis. White boxes with sold outline indicate required input from the user. White boxes with dotted outline are output from SPQ or other analysis output that can be generated from SPQ outputs.

## Results

### Assessing accuracy and functionality of SPQ

As a proof-of-principal, we tested the efficiency of SPQ by quantifying viral read counts from two published 10x scRNAseq cell culture virus infection experiments (Table 2), including two SARS-CoV-2 infection of human bronchial epithelial cells (Ravindra et al. 2021) HIV infection in a CD4 T cell model of viral infection and latency (Bradley et al. 2018). Each of these experiments reported detection of viral reads in cells within the infected cultures using CellRanger (Ravindra et al. 2021)^-^(Bradley et al. 2018). We ran both CellRanger and SPQ (using bbmap and bowite2 with default alignment parameters) on all samples from both experiments, and directly compared the count output results from SPQ to the CellRanger output.

**Table 2:**
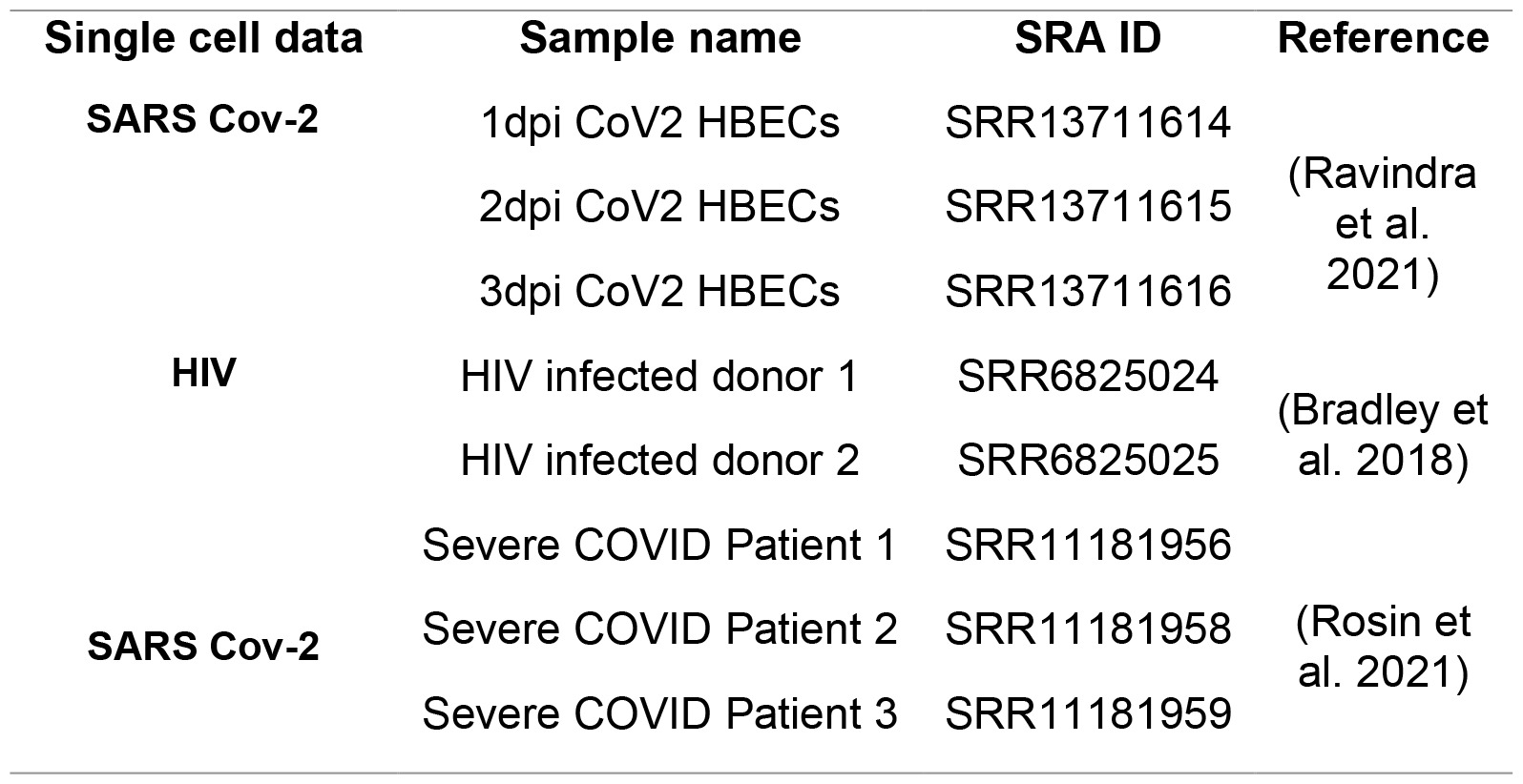
Sequencing sets used in analyses.

### HIV comparison

The single HIV infection scRNAseq study evaluated viral infection and latency in primary CD4+ T cells isolated from two healthy human donors and infected ex vivo using the recombinant pNL4-3-Δ6-drEGFP HIV strain (Bradley et al. 2018). We utilized the data set from this study because expression of enhanced green fluorescent protein (EGFP) from the engineered viral genome could be monitored to further confirm viral-mediated gene expression in cells as in internal control. We aligned reads from the donor samples infected with HIV to the human genome GRCh38 integrated with the EGFP gene using CellRanger v6.1.2. To obtain viral read alignments and compare SPQ to current methods, we performed 3 different analyses: 1) computational integration of the HIV genome into the human genome followed by aligning reads using CellRanger, which is the current method of aligning viral reads, 2) aligned reads to the human genome and then aligned viral reads to the HIV genome using SPQ with bbmap and 3) aligned viral reads to the HIV genome using SPQ with bowtie2. In both HIV-infected donor samples, we observed 100% overlap in the cells that had both EGFP and HIV reads (EGFP+HIV+) across all 3 alignment methods. The number of EGFP+HIV+ cells were 3,653 and 183 for donor samples 1 and 2, respectively. The read counts for HIV were correlated (Kendall correlation) in EGFP+HIV+ cells between methods to ensure SPQ was obtaining comparable counts to CellRanger (**Fig 2A)**. We observed high correlation of counts between CellRanger and SPQ in EGFP+HIV+ cells, showing the accuracy of both alignment methods in SPQ.

**Figure 2:**
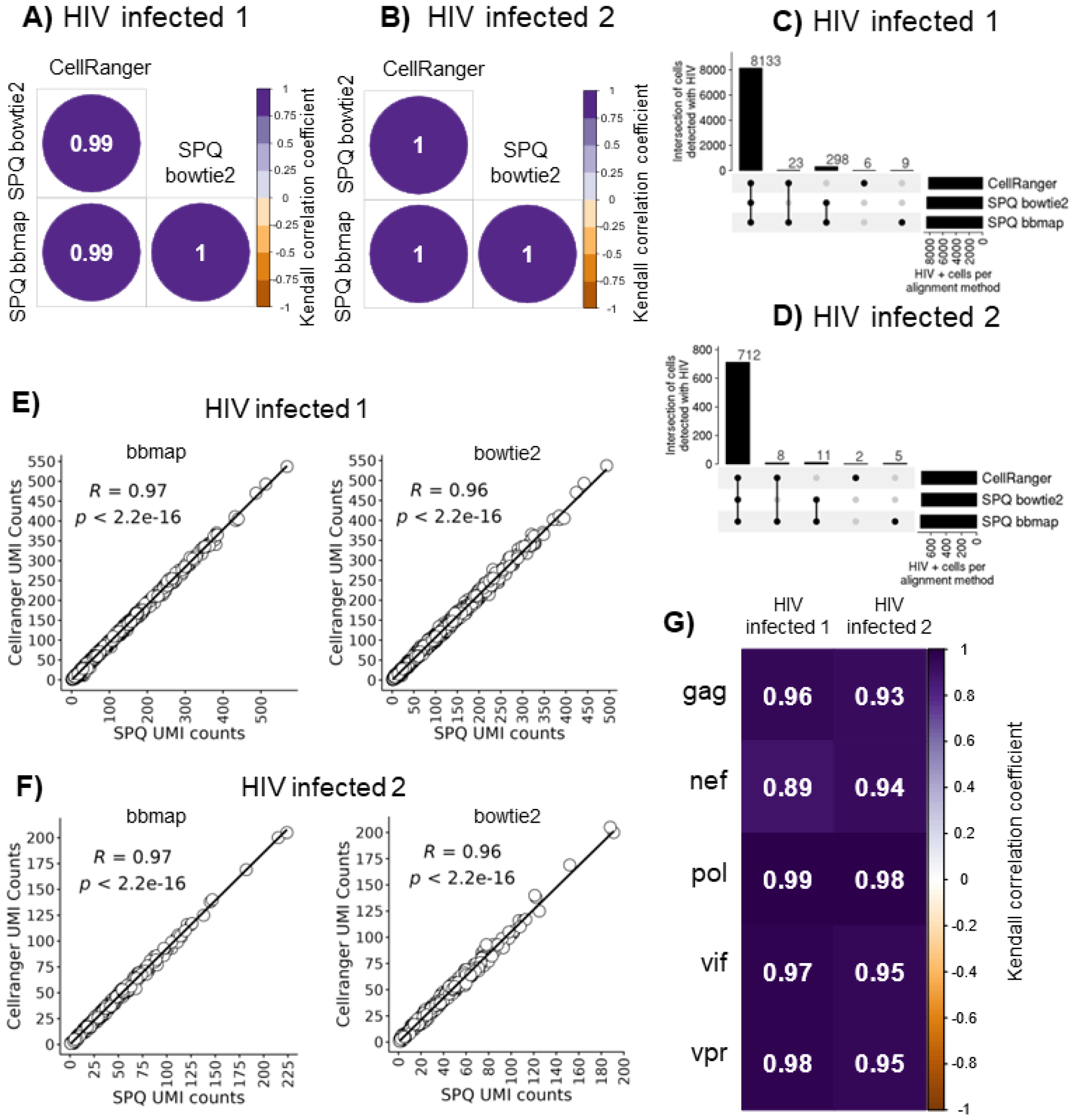
SQV comparison results from HIV-infected cells. Kendall correlation values between SPQ and CellRanger UMI counts for EGFP+HIV+ cells from **A)** HIV infected donor 1, and **B)** HIV infected donor 2. Upset plots of the overlap of EGFP-HIV+ cells detected by CellRanger and SPQ with bbmap or bowtie2 are shown for **C)** HIV infected donor 1, and **D)** HIV infected donor 2. Scatter plots showing the correlation of UMI counts in EGFP-HIV+ cells between CellRanger and SPQ with either bbmap or bowtie2 are shown for **E)** HIV infected donor 1, and **F)** HIV infected donor 2. **G)** Kendall correlation values between SPQ bbmap and bowtie2 for viral gene UMI counts for HIV-infected donor samples 1 and 2 are shown.

In addition to EGFP+HIV+ cells, both CellRanger and SPQ detected other cells with HIV reads but no EGFP reads (EGFP-HIV+). A total of 8,133 and 712 cells in HIV infected donor samples 1 and 2, respectively, were detected to be EGFP-HIV+ by all 3 alignment methods. A smaller number of cells were detected to be HIV+ by CellRanger & SPQ bbmap, SPQ bowtie2 & bbmap, or uniquely by CellRanger or SPQ bbmap (Figs 2C and D). Variations in the cells detected with viral reads were due to different alignment algorithms with different acceptable default rates of mismatches and INDELs (Table 1). We note that bbmap and CellRanger are splice aware aligners in that splice aware aligners predict where transcripts are being post transcriptionally spliced together across introns by integrating known splice junction sequence information when aligning reads to a genome. In contrast, splice unaware aligners do not account for splicing when aligning reads to genomes. We found that the HIV UMI counts in EGFP-HIV+ cells between all 3 aligners were highly correlative (Figs 2E and F), with bbmap and CellRanger UMI counts being slightly more strongly correlated since both alignment algorithms are splice aware. Lastly, we compared UMI counts for individual HIV genes (gag, nef, pol, vif and vpr) between bowtie2 and bbmap in EGFP-HIV+ cells by performing a correlation of the counts for each gene. Significant and high correlation values were observed for HIV genes in each sample (Fig 2G), showing a consistent quantification of HIV gene counts between the two aligner options in SPQ. EGFP-HIV+ read output in the HIV latency model is appropriate and expected, as viral latency is linked with selective and minimal HIV gene expression(Battistini and Sgarbanti 2014, 1).

### SARS-CoV-2 comparison

The recent COVID-19 pandemic exposed a high need for viral quantification pipelines for scRNAseq data set analyses (Liu et al. 2022). Therefore, we also tested SPQ performance on SARS-CoV-2 infection data sets. We selected a 10x scRNAseq data set of human bronchial epithelial cells (HBECs) infected with SARS-CoV-2 with time point analysis of samples collected following 1, 2, or 3 days post-infection (dpi) (Ravindra et al. 2021). Using CellRanger we aligned sequence reads first to the human genome GRCh38 followed by alignment of remaining reads to the SARS-CoV-2 genome (GCF_009858895.2) using the 3 alignment methods described above for the HIV analyses.

Whereas the SARS-CoV-2 dataset did not have an internal positive control for viral expression, all 3 alignment methods were able to identify SARS-CoV-2 sequence reads in the same 4,279, 13,998 and 26,935 cells for samples from 1,2 and 3 dpi, respectively (**Fig 3A-C**). The UMI counts in these cells were found to be highly correlated to the CellRanger UMI counts for both SPQ bbmap and bowtie2 wherein a slightly higher correlation was observed between SPQ bbmap and CellRanger with this analysis (**Figs 3D-I**). Similar to the findings with the HIV infection dataset, a smaller percentage of cells had SARS-CoV-2 detected by CellRanger & SPQ bbmap, SPQ bowtie2 & bbmap, or uniquely by CellRanger or SPQ bbmap. Lastly, we performed correlation between SPQ bowtie2 and bbmap on the quantification of UMIs for the ten SARS-CoV-2 genes. These analyses revealed significant and high correlations of the quantification of the SARS-CoV-2 genes in each sample between the two alignment methods for SPQ (**Fig 3J**).

**Figure 3:**
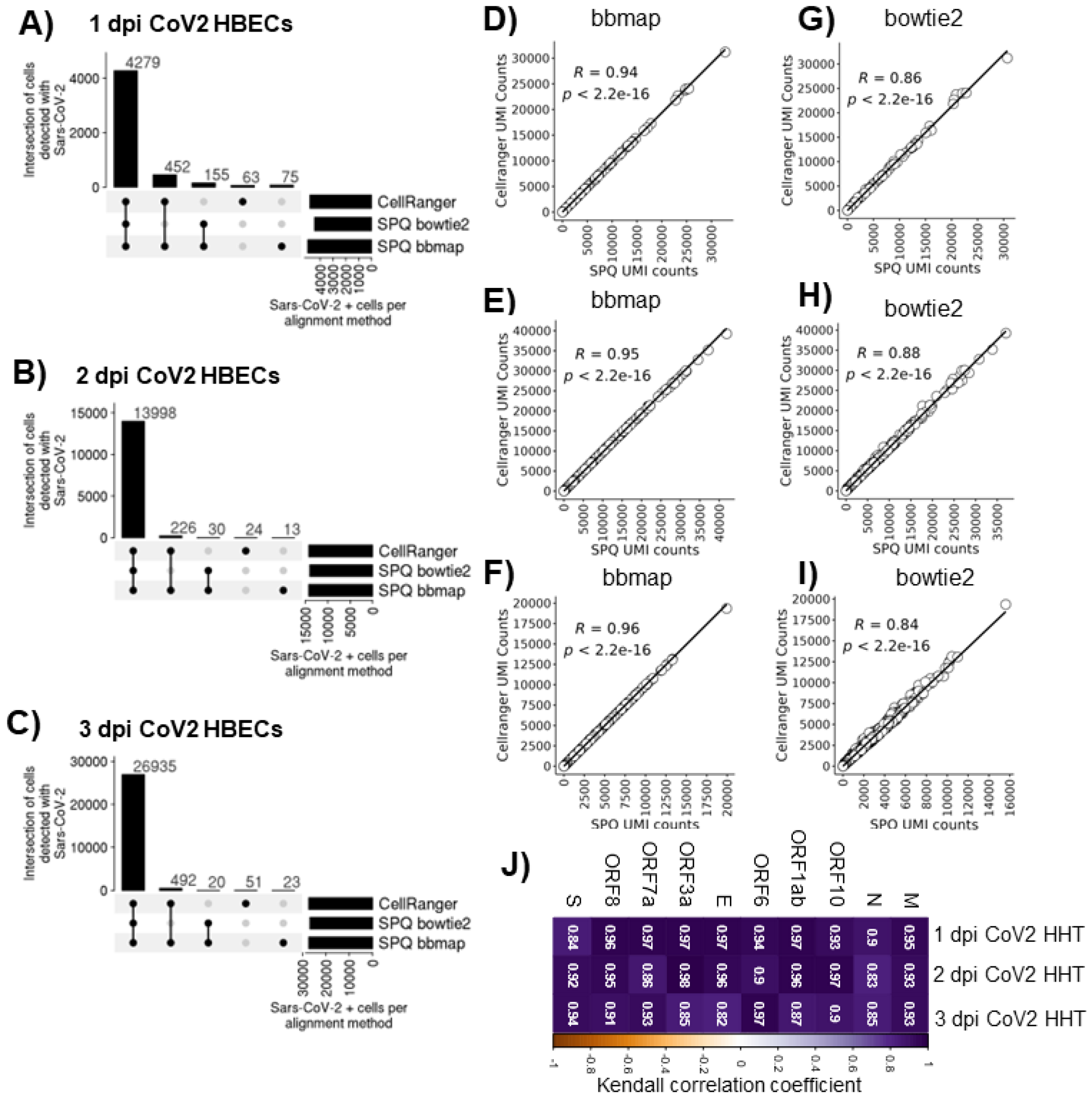
SPQ comparison results from SARS-CoV-2-infected cells. Upset plots depict the number of cells with SARS-CoV-2 RNA that overlap between analyses using CellRanger, SPQ bowtie2 and SPQ bbmap for **A)** 1dpi human bronchial epithelial cells (HBEC), **B)** 2dpi HBEC and **C)** 3dpi HBEC. Correlation of UMI counts of SARS-CoV-2 RNA+ cells using CellRanger and SPQ bbmap for **D)** 1dpi HBEC, **E)** 2dpi HBEC and **F)** 3dpi HBEC samples. Correlation of UMI counts of SARS-CoV-2 RNA+ cells using CellRanger and SPQ bowtie2 for **G)** 1dpi HBEC, **H)** 2dpi HBEC and **I)** 3dpi HBEC samples. **J)** Correlation values for SARS-CoV-2 gene UMI counts between SPQ alignment methods bbmap and bowtie2 for SARS-CoV-2-infected samples collected at 1, 2, and 3 dpi.

### Detection of co-infections

Understanding how co-infections uniquely affect the host transcriptome as well as each pathogen’s relative abundance is important for developing therapeutic interventions against infection and disease. SPQ allows for detection of multiple pathogens by providing a fasta file with all the pathogen genomes of interest. We demonstrate this ability by replicating detection of 2 viral pathogens identified in lung tissue of a patient with severe Sars-CoV-2 (Bost et al. 2020),(Rosin et al. 2021). In this study that evaluated three patients it was reported that both SARS-Cov-2 and metapneumovirus were found in patient 1 with severe COVID symptoms (Bost et al. 2020). SPQ was able to identify both viruses present in patient 1. The number of infected cells and viral read abundance of each virus is shown in **(Fig 4A-B)** with genome read coverage of each virus shown in **(Fig 4C-D)**. Only SARS-CoV-2 was found in severe COVID patients 2 and 3 by SPQ, consistent with that was previously reported (Bost et al. 2020).

**Figure 4:**
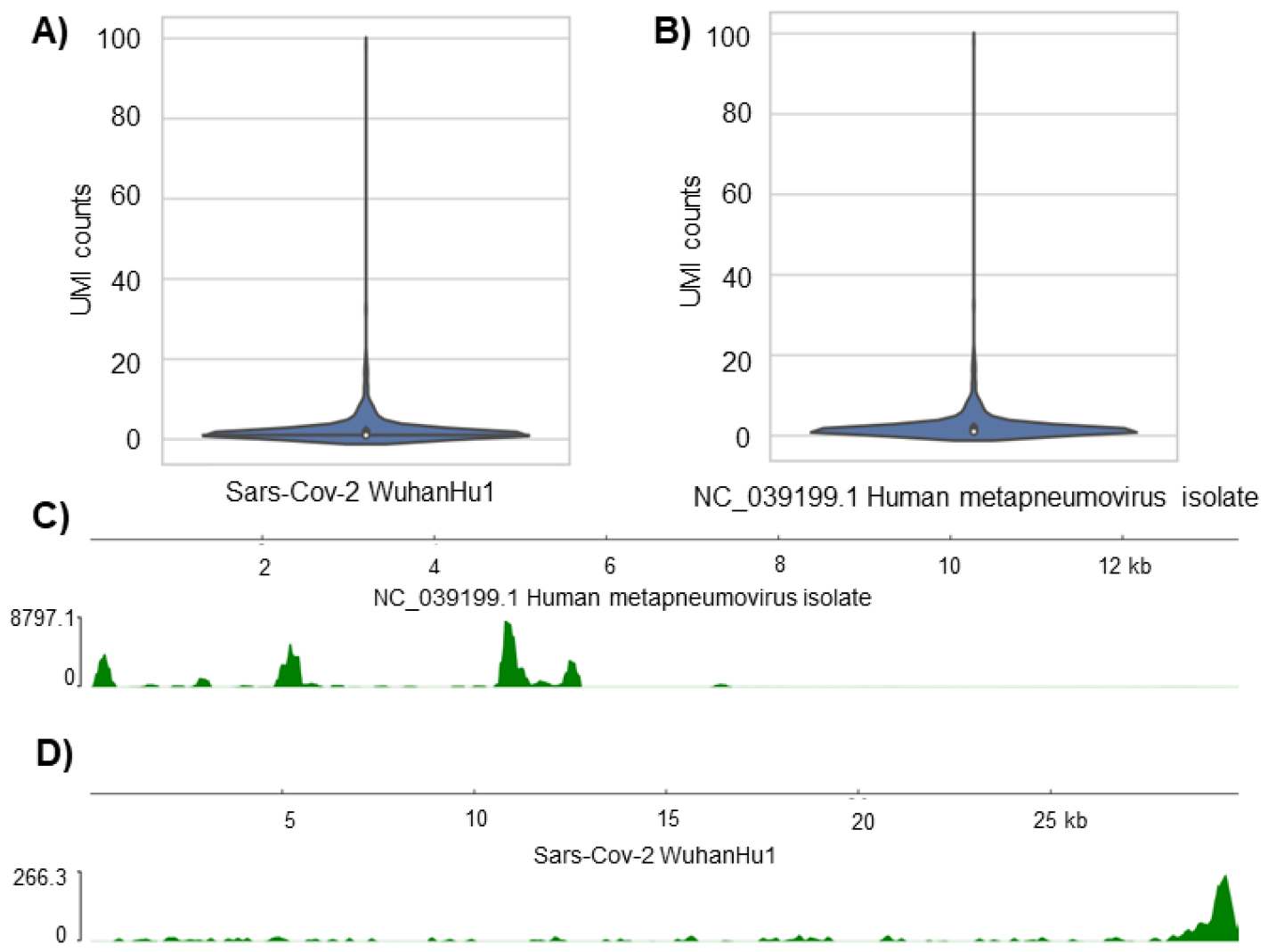
SPQ detection of co-infection. SPQ generated violin plots showing the distribution UMI counts in cells infected with Sars-CoV-2 **A)** and metapneumovirus **B)**. SPQ coverage maps of metapnemovirus **C)** and Sars-CoV-2 **D)**.

### Benchmarking SPQ against scRNAseq pathogen detection tools

Other tools used to detect pathogen derived reads in scRNAseq data include ViralTrack(Bost et al. 2020), PathogenTrack(Zhang et al. 2022) and Venus (Lee et al. 2022) all of which use STAR solo for pathogen alignment. We compared SPQ, using the bbmap aligner as it is splice aware like STAR, to these 3 other tools using HIV infected donor 2 from the previously published study (Bradley et al. 2018) (**Table 2**), examining differences in the number of HIV positive cells, abundance of HIV and CPU cost/time of each tool. There was a 81.3% overlap between all 4 tools for identifying HIV-infected cells, showing consistent results between all tools (**Fig 5A**)(Venny 2.1.0). Taking the 672 cells that were HIV positive in all 4 methods, we compared HIV abundance determined by each of the 3 tools to HIV read abundance defined by SPQ. We observe that HIV read abundance was also well correlated between SPQ and all 3 other tools (**Fig 5B-D**) with ViralTrack being the most closely in performance output to SPQ. HIV read counts from the Venus tool were higher compared to SPQ because the Venus tool counts mapped read instead of UMIs to the HIV genome but the output was still well-correlated with SPQ. Finally, we measured the computational time required for each quantification step (scpathoquant for SPQ, Viral_Track_scanning.R for ViralTrack, PathogenTrack.py count for PathogenTrack and module_detection.py for Venus) of each tool using the time command. SPQ was the fastest requiring the least amount of CPU time followed by ViralTrack, PathogenTrack and Venus which was the most computationally intensive (**Fig 5E**).

**Figure 5:**
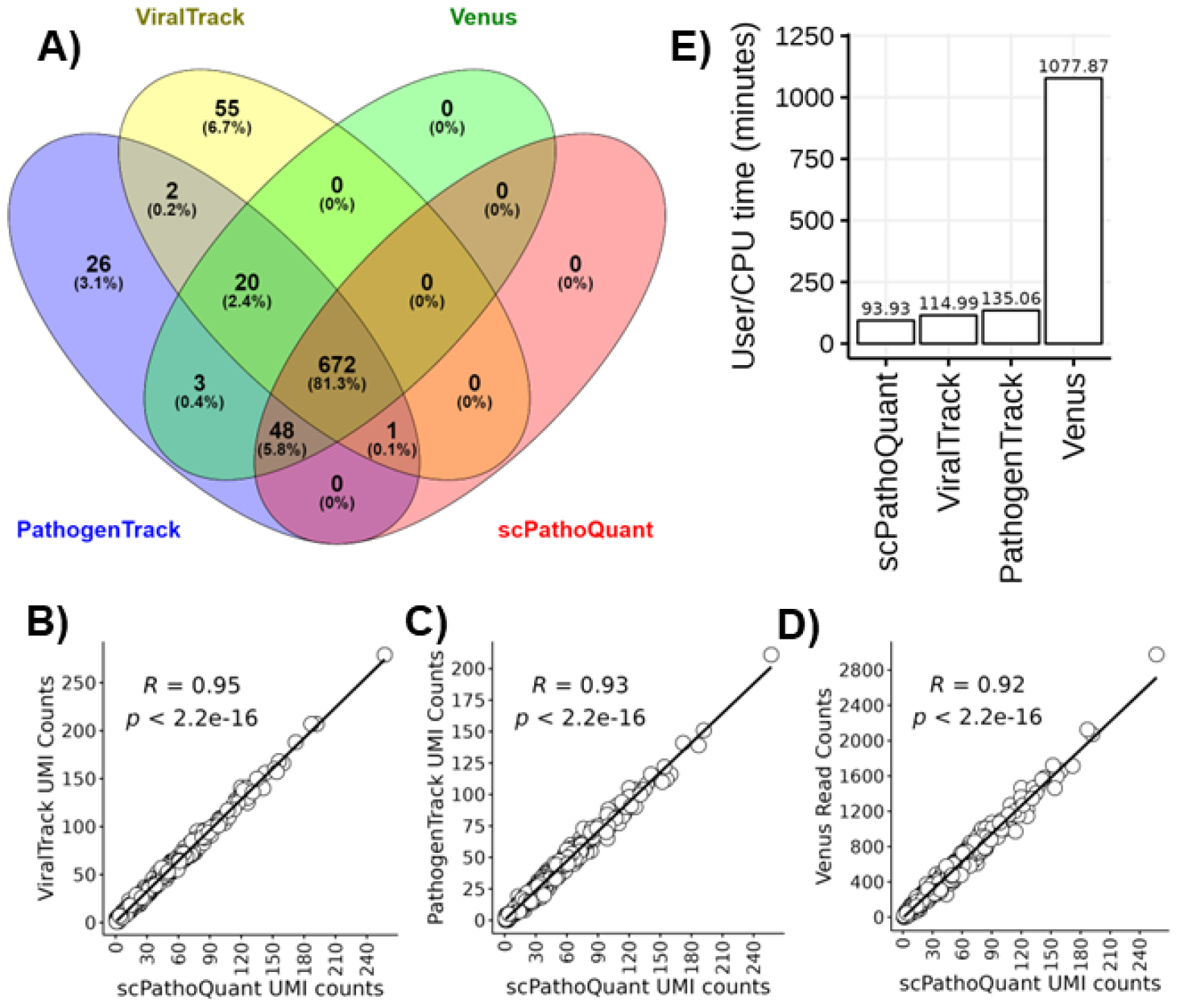
Comparison of SPQ against other viral detection tools. Venn diagram showing the overlap of HIV positive cells detected by ViralTrack, PathogenTrack, Venus and scPathoQuant **A)**. Correlation (Kendall) between HIV UMI counts of ViralTrack, PathogenTrack to scPathoQuant **B-C)** and correlation (Kendall) of read counts from Venus to UMI counts from scPathoQuant **D)**. BarPlot showing CPU/User time of each tools alignment and quantification steps **E)**.

### Availability and Future Directions

SPQ is a new bioinformatics analysis tool that provides an efficient method for quantifying pathogen genome-specific and host sequence reads in scRNAseq experiments. In addition to the three experiment examples this study, SPQ is widely applicable to analysis of any scRNAseq experiment of pathogen-infected cells. Uniquely to SPQ, individual and multiple pathogen gene expression quantification is possible, providing the user with simultaneous pathogen and host data output to quantify pathogen and host gene expression. While only viral infections were used in our examples to demonstrate SPQ applications, this tool can also be used to align and quantify sequence reads across other pathogens such as intracellular bacteria. In each case the user would provide the pathogen genome reference sequence for read mapping. These features, along with immediate figure generation, provide files that can be utilized for further downstream phylogenetic analysis. SPQ output also integrates pathogen counts into matrix and feature files from 10x data sets to allow easy and quick comprehensive analysis of pathogen/host dynamics on the single cell level. The SPQ package is available software accessible at https://github.com/galelab/scPathoQuant (DOI 10.5281/zenodo.10463670) with test codes and data sets available https://github.com/galelab/Whitmore_scPathoQuant_testSets (DOI 10.5281/zenodo.10463677) to serve as a resource for the community.

